# Calorie Restriction Maintains Mitochondrial Function and Redox Balance Avoiding Lipidomic Reprogramming during Isoproterenol-Induced Cardiac Hypertrophy

**DOI:** 10.1101/2021.07.13.452233

**Authors:** Cícera Edna Barbosa David, Aline Maria Brito Lucas, Pedro Lourenzo Oliveira Cunha, Yuana Ivia Ponte Viana, Marcos Yukio Yoshinaga, Sayuri Miyamoto, Adriano Brito Chaves Filho, Anna Lídia Nunes Varela, Alicia Juliana Kowaltowski, Heberty Tarso Facundo

## Abstract

Cardiac hypertrophy induces a metabolic shift, leading to a preferential consumption of glucose (over fatty acids) to support the high energetic demand. Typically, health cardiac tissue utilizes more fat than any other organ. Calorie restriction is a dietary procedure that induces health benefits and lifespan extension in many organisms. Given the beneficial effects of calorie restriction and the metabolic dysregulation seen during cardiac hypertrophy, we hypothesized that calorie restriction prevents cardiac hypertrophy, lipid, mitochondrial, and redox dysregulations. Strikingly, calorie restriction reversed isoproterenol-induced cardiac hypertrophy, lowered succinate driven mitochondrial H_2_O_2_ production, improved mitochondrial function (indicated as a higher Respiratory Control Ratio – RCR) and avoided mitochondrial superoxide dismutase (MnSOD) and glutathione peroxidase (GPX) repression. To gain insight into how calorie restriction could interfere with the metabolic changes induced by cardiac hypertrophy, we performed lipidomic profiling. Calorie restriction protected against the consumption of several triglycerides (TG) linked to unsaturated fatty acids, and the accumulation of TGs containing saturated fatty acids observed in hypertrophic samples. Cardiac hypertrophy induced an increase in ceramides, phosphoethanolamines and acylcarnitines (12:0, 14:0, 16:0 and 18:0) that were also reversed by calorie restriction. Altogether, our data demonstrate that hypertrophy changes the cardiac lipidome, causes mitochondrial disturbances and oxidative stress. All these changes are prevented by calorie restriction intervention *in vivo*. This study uncovers calorie restriction as a resource protect cardiac tissue and prevent cardiac hypertrophy-induced lipidomic remodeling.

## 1. Introduction

The healthy adult heart preferentially uses fatty acids (FA) to continuously derive most of its energetic demands [1]. This preferential use is shifted to less efficient glucose oxidation during pathological cardiac hypertrophy [2,3]. Although cardiac hypertrophy is adaptive, if sustained it will lead to heart failure. The transition from cardiac hypertrophy to heart failure is accompanied by numerous intracellular alterations such as reactivation of fetal genetic programing, mitochondrial dysregulation, high reactive oxygen species (ROS) production coupled to suppression in antioxidant systems, and changes in contractile proteins (for review see [4]). Despite some advancements in understanding the basic science related to cardiac hypertrophy/heart failure and recent progress in medical therapies aimed at treating heart failure, this condition persists as a leading cause of death and morbidity worldwide [5].

The cardiac tissue uses lipids acquired from circulating nonesterified free fatty acids (FFA) and esterified FA in lipoproteins [6]. Once inside cardiomyocytes acyltransferases and phosphatases (in the mitochondrial membrane and sarcoplasmic reticulum) synthesize triglycerides (TGs) and pack them into lipid droplets [7]. Endogenous TGs in lipid droplets are a dynamic depot of active substrates for ATP synthesis and a source of metabolic signaling regulators [8,9]. Increased TG synthesis (by diacylglycerol acyltransferase 1 overexpression) leads to the diversion of lipids from potentially toxic intermediates (such as ceramides) to the TG pool [10].

As stated above, the adult heart relies on fatty acids as a primary source of energy to produce ATP [1]. It is long understood that mobilized endogenous cardiac TGs are essential for cardiac energy production. Several conditions stimulate the mobilization of TGs and mitochondrial oxidation of TG-derived fatty acids examples are; diabetes and excessive catabolic/catecholamine hormone. The opposite occurs with the supply of exogenous TGs that inhibit cardiac TG mobilization and oxidation. [11].

In cardiac tissue, mitochondria are powerful coordinators of the energetic machinery. These multipurpose organelles generate about 95% of ATP used by the heart, modulate signaling, cell death, calcium homeostasis, and generate ROS. In cardiac hypertrophy, mitochondrial fatty acid oxidation is deficient due, in part, to the inhibition of carnitine-palmitoyl transferase-1 (CPT-1), which is the rate-limiting enzyme for the entry and oxidation of fatty acids into mitochondria [12] or due to reduced carnitine levels [13]. These metabolic derangements coupled with high mitochondrial ROS generation can be responsible for contractile dysfunction, ventricular dilation, and heart failure.

Studies have shown that calorie restriction, besides increasing median and maximum lifespan in mice and avoiding redox damage, is also a cardioprotective intervention [14–18]. These may be a result of less ROS production or increased antioxidant levels. Indeed, evidence shows that calorie restriction decreases mitochondrial ROS released [19–21]. Even though that idea is somehow controversial [22] animal studies have shown that calorie restriction lowers the levels of oxidatively modified lipids [23,24], proteins [25], and DNA [26]. Strikingly, calorie restriction seems to protect cardiac mitochondrial components from ROS-induced oxidative damage [20], in parallel with lower mitochondrial ROS production by this organ [19,21].

Here, we show that short-term calorie restriction promotes protection against cardiac hypertrophy through the enhancement of antioxidant enzymes (mnSOD, GPX) levels, improvement of cardiac mitochondrial function, and decreased ROS production. Additionally, we show that calorie restriction avoids the hypertrophic maladaptive lipidome. More specifically, this report implicates that this dietetic approach is capable of avoiding the accumulation of saturated and medium-chain fatty acids in cardiac triglycerides (TG) and the accumulation of specific classes of ceramides (CER), acylcarnitines (AC), and phosphoethanolamines. The changes in lipidomic, mitochondrial, and redox activities induced by calorie restriction may uncover new strategies aiming at functional improvements of cardiac hypertrophy/heart failure phenotype.

## 2. Materials and Methods

### 2.1. Animals

All protocols used in this paper were approved by the institutional Universidade Federal do Cariri’s Animal Experimentation Ethics Committee in compliance with the Guide for the Care and Use of Laboratory Animals published by the National Institutes of Health.

### 2.2. Lipid extraction

To extract lipids from homogenized cardiac samples we used the method established by Yoshida et al. [27]. Cardiac samples were homogenized in a polytron tissue grinder in icecold 10 mM phosphate buffer (pH 7.4) containing deferoxamine mesylate 100 μM. To extract lipids, we mixed the above homogenate with ice-cold methanol in PBS. Importantly, internal standards were added to the samples according to Chaves-Filho et al [28]. Then the mixture was thoroughly vortexed for 30 seconds for lipid extraction with chloroform/ethyl acetate (4:1). After centrifugation at 1500 g for 2 min at 4 °C, the lower phase containing the total lipid extract (TLE) was transferred to a new tube and dried under N2 gas. Yeast samples (10 mg) were submitted to the same procedure and used as quality controls for reproducibility analysis. Dried TLE was redissolved in isopropanol (100 μL) and injected (1 μL). Blanks and quality controls were injected every 5 and 10 samples, respectively.

### 2.3. Lipidomic analysis

We used ESI-Q-TOFMS (Triple TOF^®^ 6600, Sciex, Concord, US) interfaced with an ultra-high performance liquid chromatography (UHPLC Nexera, Shimadzu, Kyoto, Japan) to analyze TLEs. CORTECS^®^ (UPLC^®^ C18 column, 1.6 μm, 2.1 mm i.d. × 100 mm) was set to a flow rate of 0.2 mL min^-1^ and the oven temperature was maintained at 35 °C. Mobile phases A (water/acetonitrile - 60:40) and B (isopropanol/acetonitrile/water - 88:10:2) contained 10 mM ammonium acetate or 10 mM ammonium formate for experiments performed in negative or positive ionization mode, respectively. Lipids were separated by a 20 min linear gradient as follows: from 40 to 100% B over the first 10 min, hold at 100% B from 10–12 min, decreased from 100 to 40% B during 12–13 min, and hold at 40% B from 13–20 min. The MS was operated in both positive and negative ionization modes, and the scan range was set at a mass-to-charge ratio of 200–2000 Da. Information Dependent Acquisition (IDA^®^) was used to obtain data for lipid molecular species identification and quantification. We recovered the top 36 precursor ions using Analyst^®^ 1.7.1. The analysis was performed as follows: cycle time period of 1.05 s with 100 ms acquisition time for MS1 scan and 25 ms acquisition time using an ion spray voltage of −4.5 kV and 5.5 kV (for negative and positive modes, respectively) and the cone voltage at +/-80 V.

### 2.4. Lipidomics data processing

To analyze the LC-MS/MS data we manually identified lipid molecular species based on their exact masses, specific fragments, and/or neutral losses using PeakView^®^. Additionally, we used an in-house manufactured Excel-based macro. Importantly, a maximum error of 5 mDa was defined for the attribution of the precursor ion. Using MultiQuant^®^ we obtained the area of lipid species by MS data. The area ratio of each lipid was calculated by dividing the peak area of the lipid by the corresponding internal standard or using external calibration. Also, accurate area determination and correct peak detection were assured for each peak integration. The total concentration of each lipid was expressed in nmol/g of tissue and calculated by either multiplying the area ratio by the concentration of the corresponding internal standard or by external calibration curves relative to lysophosphatidylcholine (17:0). Data are presented as mean ± standard error of the mean (SEM). Data reproducibility analysis was performed by quality controls in both negative and positive ion modes. The peak area and retention time of selected lipids in quality controls (extracted from a yeast sample) were measured at the beginning, after every 10 samples, and at the end of the LC-MS/MS experiments.

### 2.5. Calorie restriction

All mice used were 60-day-old Swiss Male weighing between 25-30 g fed with a standard rodent diet ad libitum. Calorie restriction was induced by randomly dividing animals into 2 groups: mice fed ad libitum (free access to food – group control) and mice on calorie restriction of 40% of the ad libitum energy intake (Calorie Restriction group - CR). Mice were submitted to the protocol for 3 weeks and then were randomly divided into 4 other groups and treated with saline or isoproterenol (30 mg/kg per day to induce cardiac hypertrophy) [19].

### 2.6. Hypertrophy Induction

To induce cardiac hypertrophy, we treated mice daily with intraperitoneal (i.p.) injections of isoproterenol (30 mg/kg/day – ISO group). Saline (0.9% - control group) was injected in the control group. Mice fed *ad libitum* (control group) were divided into 2 subgroups: one receiving saline (*Ad libitum* control) and another receiving isoproterenol (30 mg/kg per day – *Ad libitum* ISO) for 8 days. Calorie restricted mice were also divided into 2 subgroups: one that received saline (restricted control) and another receiving isoproterenol (30 mg/kg per day – calorie-restricted ISO) for 8 days.

### 2.7. Mitochondrial Isolation

Mitochondria were isolated by differential centrifugation. Briefly, cardiac tissue was rapidly removed from the mouse and washed in a buffer containing 300 mM sucrose, 10 mM K^+^ HEPES buffer, pH 7.2, and 1 mM K^+^ EGTA at 4 °C. Then, cardiac tissue was incubated in the presence of protease Type I (Sigma-Aldrich) for 10 min. The same buffer (with 1 mg/mL BSA) was used to remove the excess of protease. The tissue was manually homogenized. Nuclei and cellular residues were pelleted by centrifugation at 1200 g for 5 min. One aliquot of this homogenate was used for western blot detection assays. To obtain the mitochondrial pellet the supernatant was recentrifuged at 9400 g for 10 min. The pellet was resuspended in a minimal amount of buffer (100 μL) and used at 100 mg/ml for mitochondrial function determination and H_2_O_2_ levels.

### 2.8. Mitochondrial Respiration

Mitochondria (100 mg protein) oxygen consumption was determined using a Clark-type electrode connected to a computer data acquisition system (Oxygraph-Hansatech, UK) in a buffer containing 150 mM KCl, 10 mM HEPES, 2 mM MgCl_2_, and 2 mM KH_2_PO_4_ pH 7.2 (KOH). To induce state 2, we added 4 mM succinate. Then, we added 1 mM ADP to induce state 3 and 1 μg/ml oligomycin was added to induce state 4. The Respiratory Control Ratio (RCR) was calculated as the ratio of the slope of State 3 (induced by ADP)/slope State 4 (induced by oligomycin). After each run, mitochondria (in state 4) were used to measure H_2_O_2_ production as described below.

### 2.9. H_2_O_2_ Measurements

To measure mitochondrial H_2_O_2_ production, cardiac mitochondria in state 4 (in the presence of 1 μg/mL oligomycin) were incubated (protected from light) with Amplex red (50 μmol/L) and horseradish peroxidase (1 U/mL) at 37°C for 60 min, pH 7.2. Respiration buffer consisted in 150 mM KCl, 10 mM HEPES, 2 mM MgCl_2_, 2 mM KH_2_PO_4_, 4 mM succinate, and 1 μg/mL oligomycin, pH 7.2 (KOH). After an incubation period, tubes were centrifuged at 9300 g for 2 minutes, the supernatant was transferred to a cuvette, and the absorbance measured at 560 nm. Background absorbance was determined by incubating Amplex red and horseradish peroxidase without the sample. H_2_O_2_ released was calculated in μmol/mg protein using a calibration curve created using H_2_O_2_ standards.

### 2.10. Western blots

Proteins were harvested from mouse cardiac tissues and subjected to different treatments, as described above. Protein was submitted to electrophoresis in SDS-PAGE (4-10%) and transferred to PVDF membranes. The membrane blot was blocked (room temperature) using Tris-buffered saline pH 7.5 (TBS) containing nonfat milk (5%). Primary antibodies were rabbit anti-SOD2/MnSOD (Abcam – 1:1000) and Rabbit monoclonal [EPR3312] to Glutathione Peroxidase 1 (ABCAM – 1:2000) at 4 °C, overnight. Membrane blots were washed 3 times with TBS containing Tween-20 (TBS-T; 0.1%), blocked for 1 hour, and probed with IRDye^®^ 800CW Goat anti-Rabbit IgG Secondary Antibody (LI-COR - 1:20000). After washing four times with TBS-T, green fluorescence was detected using an Odyssey detection device. Densitometry was performed using nonsaturated exposed fluorescent membranes and quantified using Image J software. To confirm the linear range of the signal, we performed multiple exposures for every experiment. Levels of proteins in each lane were normalized to loading protein content (detected by Ponceau stain) and expressed relative to control (set as 100%).

### 2.11. Protein Content

Protein content in each sample was estimated using the Bradford method, with bovine serum albumin as a standard.

### 2.12. Statistical Analysis

Statistical analysis. Lipidomics statistical analyses was performed with Metaboanalyst (website: www.metaboanalyst.ca). All data were log transformed prior to statistical analyses. Briefly, all groups were compared by multivariate analysis (sPLS-DA). We also performed heatmap plots using PLS-DA VIP. For pairwise comparisons, we performed volcano plots assuming unequal variances and multivariate analysis (OPLS-DA), p-value threshold < 0.05 and fold change set to >1.5. Graphics were generated using metaboanalyst, graphPad Prism 6 and Excel. We used Metaboanalyst for analyses as described elsewhere [29]. Pearson correlation analysis between individual lipids and HW/TL was performed using Graphpad Prism. p < 0.05 was considered statistically significant. Data in Figs 1 and 4 are presented as mean ± S.E.M. Student’s t test or one-way analysis of variance followed by a Tukey’s post hoc test was performed where indicated. p < 0.05 was considered statistically significant.t.

**Fig. 1.**
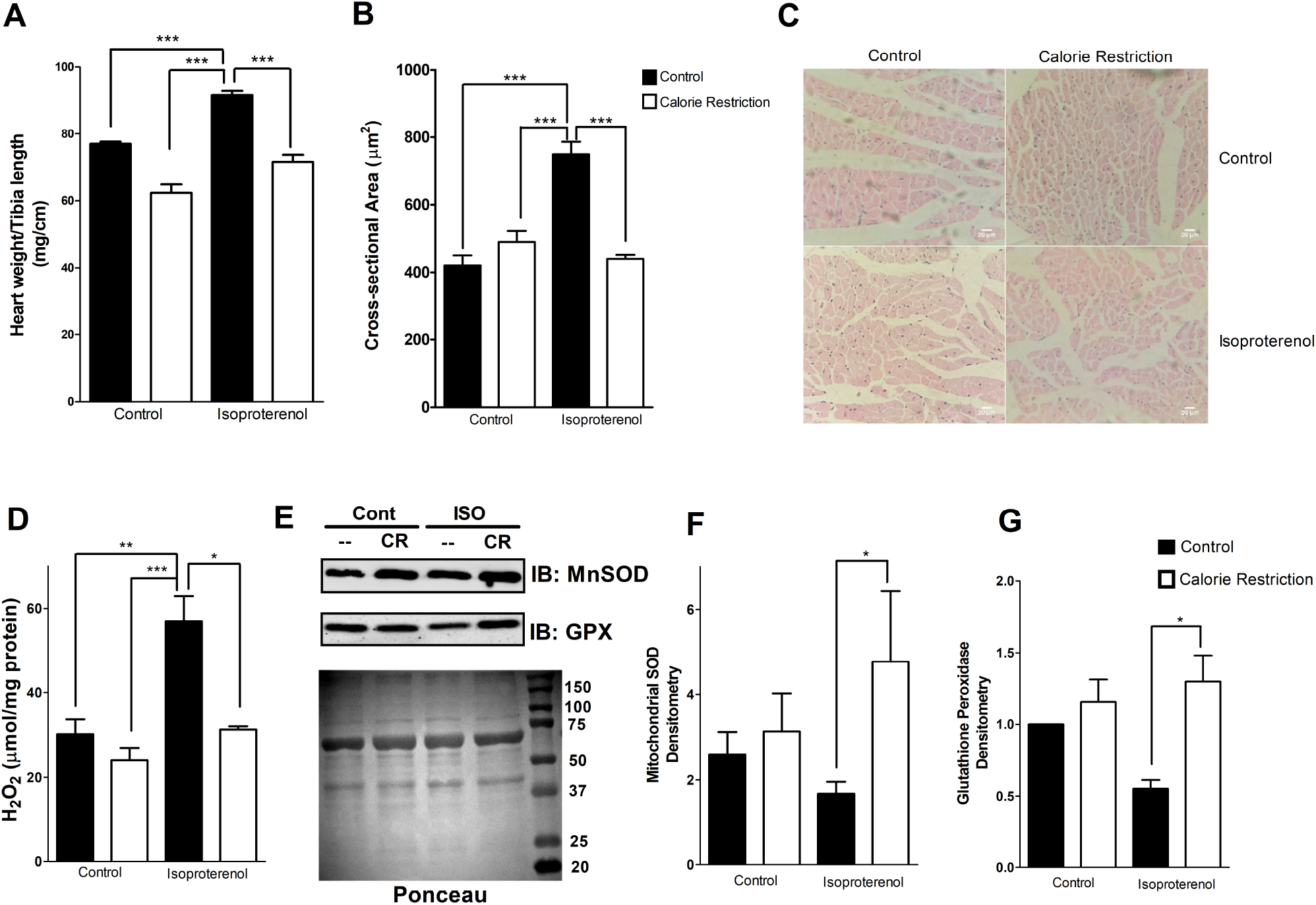
Calorie restriction avoids cardiac hypertrophy and oxidative stress. (A) Heart weight/tibia length ratio. (B) Cross-sectional area of cardiomyocytes, in mm^2^. (C) Heart sections stained with hematoxylin-eosin. (D) H_2_O_2_ production by mouse heart mitochondria (100 mg protein/mL) incubated at 37 °C in 150 mM KCl, 10 mM HEPES, 4 mM succinate, 2 mM MgCl_2_, 2 mM KH_2_PO_4_, 1 μg/mL oligomycin, pH 7.2 (KOH). Isolated mitochondria from each condition were incubated (protected from light) with Amplex red (50 μmol/L) and horseradish peroxidase (1 U/mL) for 60 min, pH 7.2. (E) Representative mitochondrial superoxide dismutase (MnSOD) and glutatione peroxidase (GPX) immunoblotting showing the effect of calorie restriction on controls and hypertrophic samples submitted or not to calorie restriction. Ponceau stain served as a loading control. Cont = control, ISO = isoproterenol-treated samples and CR = calorie restriction. (F) Quantification of 4 individual MnSOD western blots relative to ponceau stain. (G) Quantification of 4 individual GPX western blots relative to ponceau stain. Values are expressed as mean ± SE, n = at least 4 per group. * P<0.05, **P < 0.01, ***P<0.001.

## 3. Results

### 3.1. Calorie restriction blocks cardiac hypertrophy by affecting mitochondrial ROS release, antioxidant enzyme repression and improving mitochondrial function

To study the possible anti-hypertrophic effects of calorie restriction in mice, we first examined the effect of calorie restriction in control mice. Control mice submitted to 4 weeks of 40% food deprivation displayed no alteration in heart weight/tibia length (HW/TL) relationship (a gross indicator of cardiac hypertrophy - Fig 1A) and myocyte cross-sectional area (m-CSA - Fig 1B, C). This result indicates that calorie restriction *per se* does not induce any pathological cardiac hypertrophic growth. To investigate the potential anti-hypertrophic effects of calorie restriction, we injected mice with isoproterenol (ISO - 30 mg/kg/day daily) for the last week of the calorie restriction protocol (8 days). Mice fed *ad libitum* and treated with isoproterenol displayed higher HW/TL and m-CSA than controls. However, this was markedly reduced in calorie-restricted mice treated with ISO (Figs 1A, B, and C). These results suggest that calorie restriction is capable of blocking the induction of cardiac hypertrophy in mice.

To gain evidence on the beneficial effects of calorie restriction (during hypertrophy), we tested the mitochondrial ROS production (by probing for H_2_O_2_ release). Oxidative stress has been implicated in cardiac hypertrophy, particularly during the transition to heart failure. Indeed, antioxidants block cardiac hypertrophy (please see (please see [30]). Here, we decided to test H_2_O_2_ generation by mitochondria from hypertrophic samples submitted or not to calorie restriction. To force high production of H_2_O_2_, we energized isolated mitochondria with the complex II substrate succinate. This procedure allows for transfer of electrons from complex II to complex I (reverse electron transfer), generating high levels of H_2_O_2_ [31] which are sensitive to rotenone (result not shown). Fig. 1D shows that mitochondria isolated from hypertrophic hearts energized as described above generate significant amounts of H_2_O_2_. This was associated with a lower respiratory control ratios (RCR, Table 1), mitochondrial superoxide dismutase levels (MnSOD, Fig. 1E, F), and glutathione peroxidase (GPX, Fig. 1E, G). Remarkably, H_2_O_2_ production was significantly reduced in calorie-restricted mitochondria (Fig. 1D). Calorie restriction significantly improved the RCR (Table 1), the levels of MnSOD (Fig 1E, F), and GPX (Fig. 1E, G). Accordingly, we previously found that caloric restriction increases heart glutathione peroxidase activity [19]. Here, we also find that glutathione peroxidase levels were significantly improved in calorie restricted mice treated with isoproterenol (Figure 1E,G). Interestingly, glutathione peroxidase activity was negatively correlated with hypertrophy (data not shown). These results confirm findings [15,16,19] indicating that calorie restriction avoids cardiac hypertrophy and impacts mitochondria and oxidative stress.

**Table 1:** Mitochondrial respiratory control ratio (RCR) of controls and calorie-restricted mice treated or not with isoproterenol.

| Mitochondrial Respiratory Control Ratio (RCR) |  |  |  |
| --- | --- | --- | --- |
| ALC | RCC | AL_i | RCC_i |
| 3.34 ± 0.39 | 3.49 ± 0.85 | 1.29 ± 0.23<sup>A,B</sup> | 4.4 ± 1.18<sup>C</sup> |

P < 0.05
A = AL\_i x ALC
B = AL\_i x CRC
C = AL\_i X CRC\_i

### 3.2. Calorie restriction induces reprogramming of the lipid profile during isoproterenol-induced cardiac hypertrophy

To understand the impact of isoproterenol-induced cardiac hypertrophy and the beneficial effects of calorie restriction on the cardiac lipidome, we conducted a multivariate partial least square - discriminant analysis (PLS-DA). The graph in Figure 2A shows a clear separation between isoproterenol-treated samples and all other groups. This finding is consistent with substantial alterations in the cardiac tissue lipidome of isoproterenol-injected mice (AL_i), and with calorie restriction preserving lipid composition relative to the non-hypertrophic group (Fig. 2A). Strikingly, a loadings PLS-DA plot shows that the spatial separation of groups is mostly explained by alterations in triacylglycerols (TG) and acylcarnitines (AC) (Fig. 2B). Volcano plot analysis revealed several altered lipid species (represented by their corresponding classes) in the isoproterenol-treated group (AL_i) relative to controls (ALC), i.e. *ad libitum* animals with or without isoproterenol (Figure 2C). Importantly, calorie-restricted mice treated (CRC_i) or not ( CRC) with isoproterenol showed a very similar lipid profile (Fig. 2D). In contrast, several lipid species were altered in the AL_i group as compared to the other groups (Fig. S2). Comparatively, the CRC_i group displayed considerably less altered lipids relative to ALC and CRC groups, again supporting that calorie restriction prevented lipidome alterations induced by ISO (Fig. S2). Similar findings were obtained by univariate analysis using the *t* test (p < 0.05) displayed by Venn diagram analysis (Fig. S3).

**Fig. 2.**
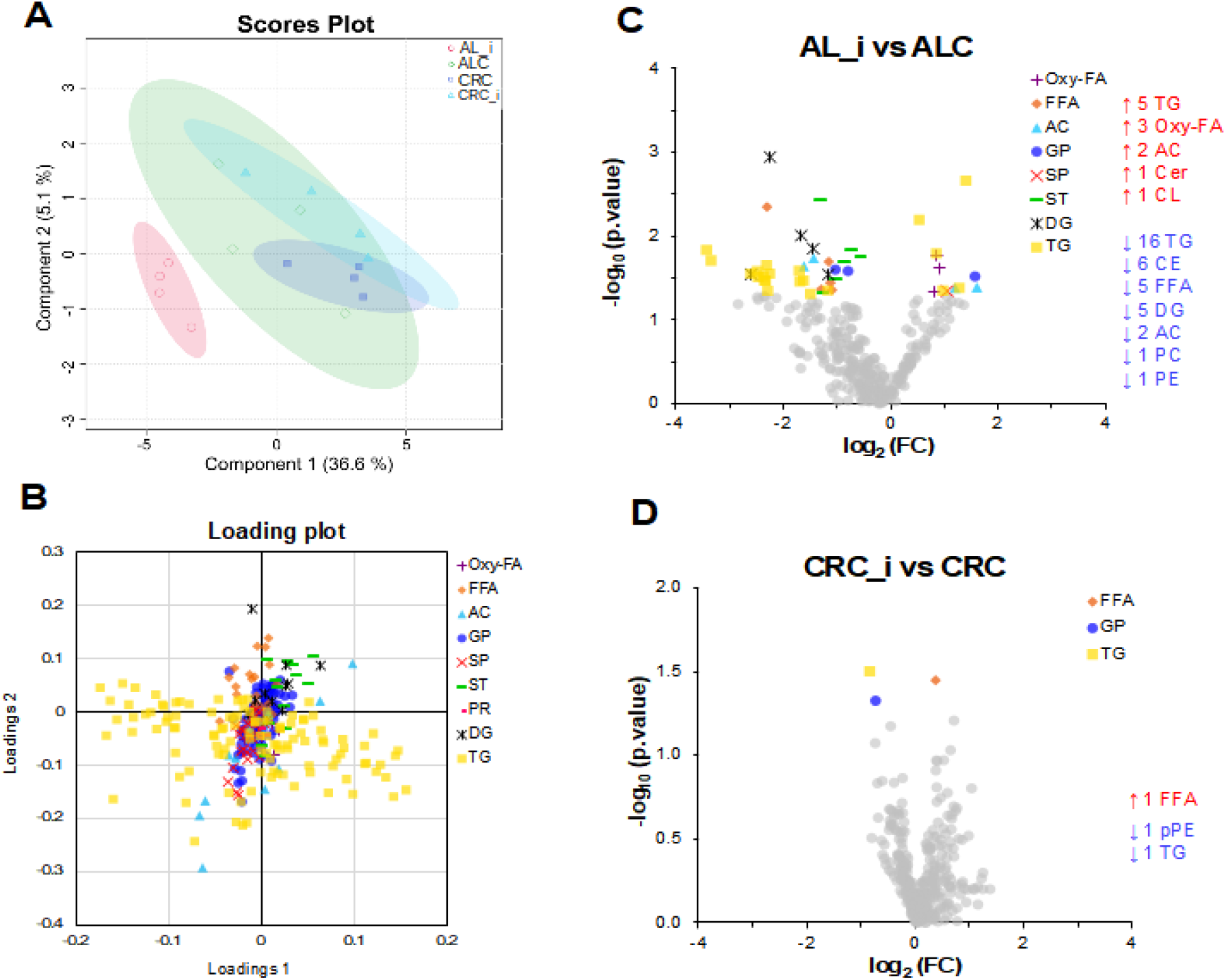
Multivariate and univariate analyses reveal drastic alterations in the cardiac tissue lipidome of mice isoproterenol-treated. Multivariate analysis was performed by partial least squares-discriminant analysis (PLS-DA). Score (A) and loading (B) plots of PLS-DA are displayed. (C) Volcano plot of the pairwise AL_i versus ALC group comparison. (D) Volcano plot of the pairwise CRC_i versus CRC group comparison. Volcano plots are represented by log2 (fold change) plotted against-log_10_ (p value). Statistical significance was evaluated by t-tests (p<0.05) and the fold change was set to >1.0. Abbreviations: Oxy-FA, oxidized fatty acid; FFA, free fatty acid; AC, acylcarnitine; GP, glycerophospholipid; SP, sphingolipid; ST, sterol lipid; PR, prenol lipid; DG, diacylglycerol; TG, triacylglycerol; Cer, ceramide; CL, cardiolipin; CE, cholesteryl ester; PC, phosphatidylcholine; PE, phosphatidylethanolamine; pPE, plasmenyl phosphatidylethanolamine.

To obtain further details about the lipidome alterations linked to cardiac tissue hypertrophy, we performed a correlation analysis. Several TG species that significantly correlated with heart weight/tibia length (HW/TL - a gross marker of cardiac hypertrophy) were TG species (Fig. 3A). This finding supports evidence that ISO induces major alterations in the TG profile of the AL_i group. Among TG species, those esterified to saturated fatty acids were positively correlated with HW/TL (Fig. 3A-B). The relative increase in abundance of TG esterified to medium-chain saturated fatty acids (12 carbons) in ISO-treated animals is intriguing, but may suggest a possible shift in metabolism of the hypertrophic cell to store (or use) a faster energy source, since medium-chain fatty acids do not need to be activated by the carnitine transport system to enter mitochondria, and are preferentially oxidized [32]. On the other hand, TG species negatively correlated with heart weight were mainly esterified to mono (MUFA) and polyunsaturated fatty acids (PUFA) (Fig. 3A,B). In addition, other lipid species such as long-chain AC, ceramides and pPE showed a positive correlation with heart weight, while a few species, including short-chain AC and linoleic acid oxidized species, displayed negative correlation with heart weight (Fig. 3C). These results suggest that cardiac hypertrophy induces a lipidomic reprograming in the cardiac tissue and that calorie restriction blocks it.

**Fig. 3.**
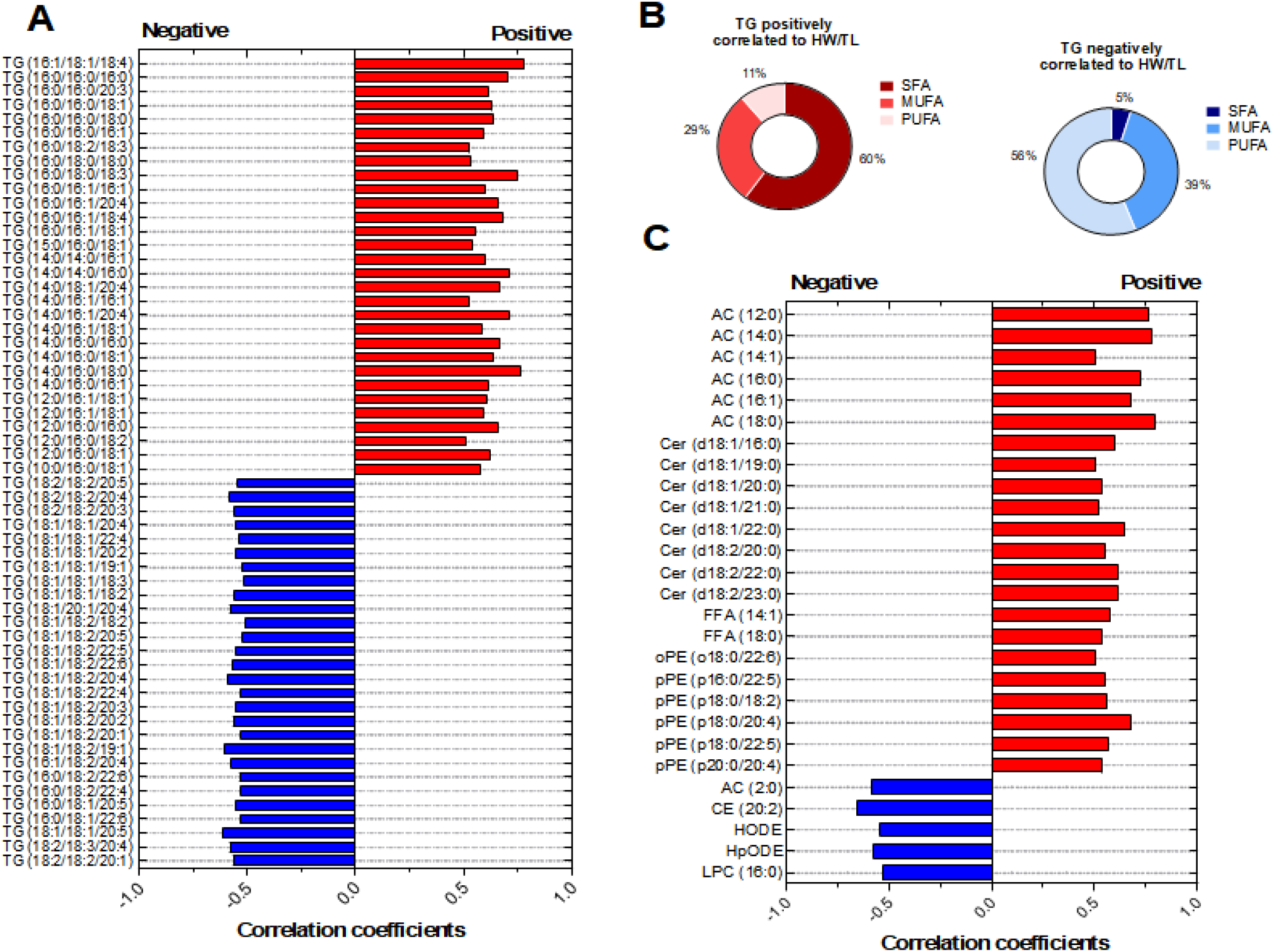
Heart weight shows significant correlation with triacylglycerols, acylcarnitines, ceramides and ether phosphatidylethanolamine. Correlation coefficients were obtained by Spearman analysis (p < 0.05). (A) TG correlated with heart weight/tibia length displayed in red (positive correlation) or blue (negative correlation). (B) Other lipid species correlated with heart weight/tibia length are displayed in red (positive correlation) or blue (negative correlation). Abbreviations: TG, triacylglycerol; AC, acylcarnitine; Cer, ceramide; FFA, free fatty acid; oPE, plasmanyl phosphatidylethanolamine; pPE, plasmenyl phosphatidylethanolamine; CE, cholesteryl ester; HODE, hydroxyoctadecadienoic acid; HpODE, hydroperoxyoctadecadienoic acid; LPC, lysophosphatidylcholine.

### 3.3. Analysis of cardiac fatty acids esterified to TG of mice subjected to different treatments

Major lipidome alterations linked to cardiac tissue hypertrophy induced by ISO were observed in TG species. Thus, we investigated the TG fatty acyl composition in all experimental groups. Major alterations were observed in the AL_i group and linked to increased relative percentage of saturated fatty acids, such as 12:0, 16:0 and 18:0, in TG, as compared to the CRC_i group. On the other hand, AL_i mice showed a significant decrease in TG esterified to mono and polyunsaturated fatty acids, such as 18:1, 18:2, 20:1, and 22:4. Notably, the CRC_i group reestablished the levels of TG linked to mono and polyunsaturated fatty acids to control levels (ALC and CRC groups) and decreased the contribution of saturated fatty acids (Figure 4).

**Fig. 4.**
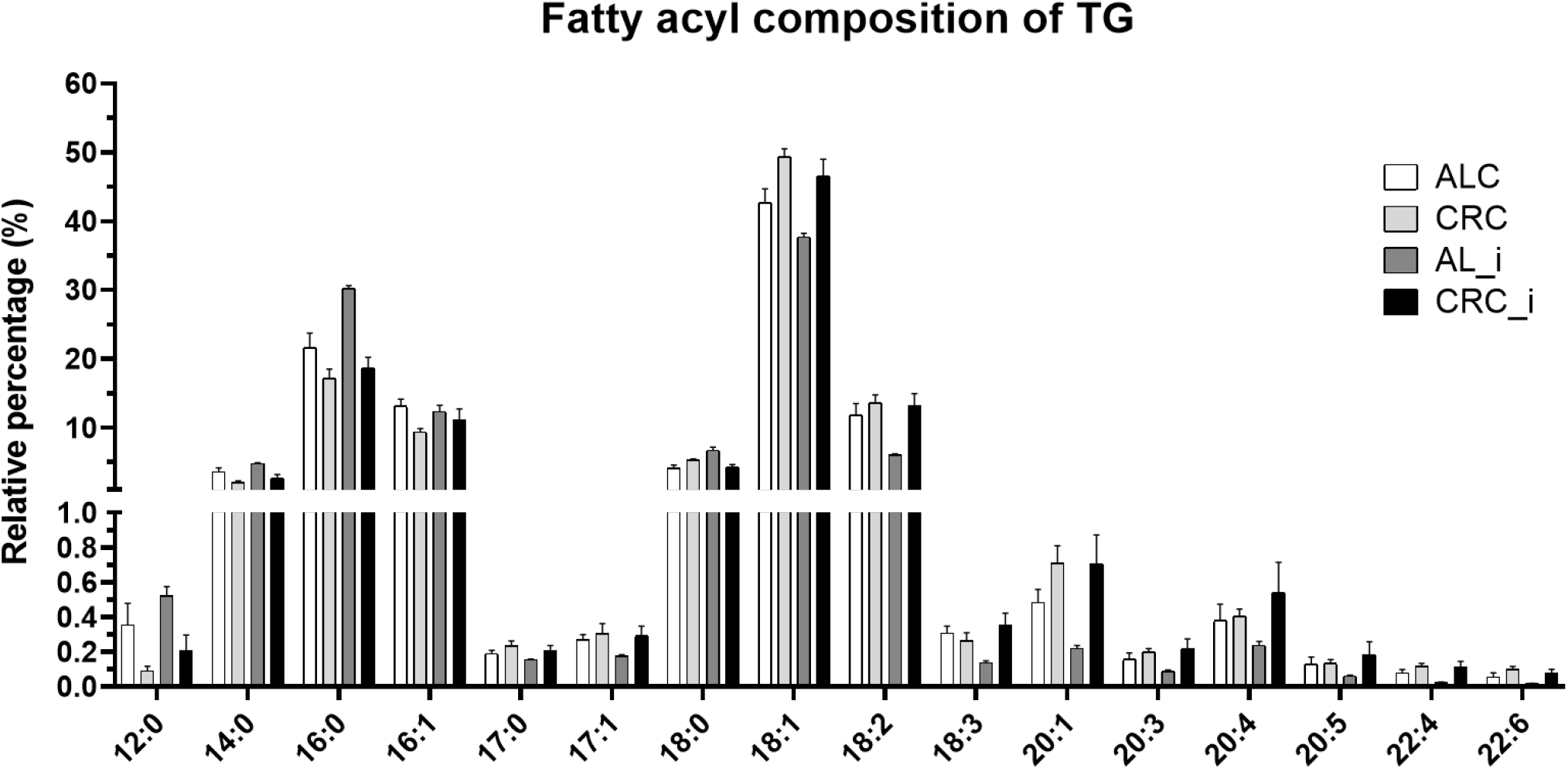
Caloric restriction avoids TG remodeling in isoproterenol-induced hypertrophic heart. Data of relative percentage from total TG are shown as mean ± standard error of mean (SEM). Only the most abundant fatty acyl chains esterified to TG are displayed. All groups were compared by one-way ANOVA followed by Tukey’s post-hoc. * P<0.05, **P < 0.01, ***P<0.001.

## 4. Discussion

The heart oxidizes more fat than any other organ in the body. This is necessary to sustain the high energetic demand of persistent contraction/relaxation, performed against a pressure load. Indeed, changes in metabolic profiles are cental in pathologic cardiac conditions such as hypertrophy. In this study, we have advanced the knowledge of mitochondrial derangements during cardiac hypertrophy, by demonstrating that mitochondria (isolated from cardiac hypertrophic samples) consume less oxygen, produce more H_2_O_2_, and develop antioxidant impairment. Cardiac hypertrophy also changes the cardiac lipidomic profile. Strikingly, we have found that calorie restriction is capable to prevent cardiac hypertrophy in mice and avoid changes in the hypertrophic lipid profile.

Together, the results bear evidence fora quantitative decrease in TG linked to linoleic acid (LA, 18:2) and other PUFA with concomitant enrichment in TG linked to saturated FAs in AL_i mice. This data suggests that 1-lipolysis of TG is induced by acute injections of ISO; 2) beta-oxidation of LA, which is one of the most beta-oxidizable fatty acids in both mitochondria and peroxisome (Refs); and 3) de novo lipogenesis of saturated FAs such as palmitic acid (16:0). The data also show that AL_i mice have cardiac accumulation of long-chain ACs, and that these positively correlate with heart weight, which may indicate impaired mitochondrial FA beta oxidation. Given that LA is largely consumed by beta-oxidation in cardiac tissues, and mitochondrial FA beta-oxidation is impaired in AL_i mice, a significant contribution of peroxisomal beta-oxidation is plausible. Alternatively, quantitative beta-oxidation of LA in mitochondria may lead to allosteric inhibition of CPT-1 by malonyl-CoA. We also find dysregulated FA beta-oxidation in AL_i mice, which is likely to stimulate de novo lipogenesis, explaining the positive correlation of TG enriched by saturated fatty acids and ceramides with heart weight. Finally, calorie restriction prevented hypertrophy, reestablished mitochondrial oxidation coupling and avoiding intense lipidome remodeling induced by isoproterenol, most notably in the pools of TGs. One possibility is that calorie restriction might counteract the lipolytic effects of ISO, but the exact mechanisms remain unclear. Strikingly, calorie restriction reverses the maladaptive accumulation of medium-chain TG fatty acids. Additionally, this dietetic intervention controls the decrease in unsaturated TGs seen in the hypertrophic profile.

From a mechanistic point of view, our data show that changes in the profile of cardiac lipids, assessed during cardiac hypertrophy, correlates with the degree of cardiac hypertrophic growth, and calorie restriction is capable of reversing this. Importantly, this dietary approach does not seem to change all lipid classes in the same way. Calorie restriction seems to target the hypertrophy-altered lipids, but does not induce significant changes compared to control samples. The lipid classes are mainly (but not limited to) TG and acylcarnitines. Conversely, a prior report shows that cardiac hypertrophy induced by transverse aortic constriction in mice induces an increase in phospholipids and sphingolipids [33]. In our model of isoproterenol-induced cardiac hypertrophy, total or individual myocardial phospholipids and sphingolipids did not change. We did find some positive correlation between cardiac hypertrophy and some classes of phosphatidylethanolamines (PE). Of note, we found that low (medium-chain) carbon number and saturated TGs were higher in hypertrophic samples. The cardiac tissue relies on long-chain fatty acids as a primary source of carbons for energy production. Additionally, some papers have pointed out that cardiac TGs are important for the storage of substrates actively used for ATP synthesis and signaling [34,35]. Cardiomyocytes are densely packed with mitochondria. They produce and consume large quantities of ATP very efficiently. On the other hand, slight changes in substrate availability/utilization directly impact cardiac energetics [36]. Cardiac energetic supply comes primarily from circulating fatty acids and from TG, which used to be considered a static, inactive fat deposit, but this point of view has recently changed. Cardiac TGs are currently considered a source of intracellular fatty acids (mainly long-chain fatty acid) [34,35]. We believe the major problem with the hypertrophic TG pool is not the total amount of TG (please see Figure S1), but their composition. Indeed, the cardiac hypertrophic TG pool not only loses long-chain fatty acid unsaturations but also accumulates saturated and medium-chain (not usual) fatty acids. Interestingly, Stegemann and collaborators have found that low carbon number and low double bond TG content in plasma from human subjects had the most consistent and strongest associations with the risk to developf cardiovascular disease [37]. This could represent a shift in metabolism to store (or use) a faster energy source as low carbon fatty acids do not need the carnitine transport system to enter mitochondria [32]. Another possibility is that hypertrophic cells accumulate low double bond TG by reflecting increased TG pool uptake and availability (however, mice had no changes in plasma TG, results not shown). An incomplete breakdown of fatty acids (poor fatty acid oxidation) is also possible. High levels of dietary long-chain monounsaturated fatty acids (in humans) are associated with a greater incidence of congestive heart failure [38]. Indeed, impaired fatty acid oxidation may contribute to intramyocardial triglyceride accumulation and cardiac contractile dysfunction in Zucker obese rats [39]. In chronic heart failure induced by pressure-overload, a decrease in endogenous TG turnover was accompanied by decreased fatty acid oxidation [34]. The TG pool could buffer the cytosolic accumulation of medium-chain/saturated fatty acids as an adaptive mechanism against cell toxicity. Finally, TG accumulation in lipid droplets may be beneficial. Indeed, lipid droplets could serve as a buffer against lipid toxicity [40].

Our study also found other enriched lipids during cardiac hypertrophy. We highlight the changes in ceramides (well-studied sphingolipids), mainly composed of 18:1 and 18:2 fatty acids. Even though the total ceramides in controls or isoproterenol-treated samples are among the less abundant lipids (less than 1 nmol/mg tissue - Figure S1), we cannot not exclude this class of lipids as important metabolic mediators. Previous studies have indicated that high levels of d18:1 and d18:2 ceramides are associated with cardiac hypertrophy [33]. Others have shown that increased total levels of ceramides are a hallmark of heart failure-induced lipid metabolism dysregulations [41,42]. Additionally, insulin resistance with inhibition of AKT signaling concomitant with abnormal AMPK activity were linked to increased myocardial levels of ceramide and PKC activation [43]. Typically, most of the cardiac TG pool is derived from de novo lipogenesis since the incorporation of TG-rich lipoprotein particles is limited in the healthy heart [6]. *De novo* lipogenesis is highly regulated and directs some lipids to FA oxidation and some to the TG pool [44]. Fatty acid oxidation dysregulations may lead to abnormal diversion of acyl-CoA intermediates to the TAG pool and lipotoxic ceramide formation. Here we show (Figure 4) that palmitic acid (16:0) accumulates in TGs of hypertrophic samples in a manner reversed by calorie restriction. Conversely, the levels of oleate (18:1) and linoleate (18:2) are downregulated in TGs from hypertrophic hearts. Strikingly, downregulation of oleate and linoleic acid in TGs from hypertrophic samples (Figure 4) was reversed by calorie restriction. Because palmitic acid is a crucial mediator of lipotoxicity, causing inflamatory signalling and ceramide upregulation [45] a possibility exists that calorie restriction improves palmitic acid levels and this influences ceramide levels. Indeed, impaired mitochondrial function found in hypertrophic tissue may be caused by higher toxic amounts of ceramides, as seen in the positive correlation of ceramides with heart weight. It is quite possible that palmitic acid accumulation in hypertrophic samples TGs is due to defective beta oxidation. Also of note, the TG reserve of oleic/linoleic acid in hypertrophic samples is impaired, which may avoid TG turnover that would rescue palmitic acid release from TG, since oleate is able to block the capacity of palmitate to induce pathological remodeling [40]. Another possibility could be a downregulation in the enzyme diacylglycerol acyltransferase 1 (DGAT1). Indeed, severe heart failure downregulates cardiac DGAT1 mRNA levels [41]. An open question here would be what are the DGAT1 levels after calorie restriction in mice, since calorie restriction corrected TG levels of palmitic, oleic and linoleic acid. This must be evaluated in future studies.

Cardiac functional and structural remodeling during cardiac hypertrophy transition to heart failure is preceded by metabolic changes that trigger myocardial dysfunction [45,46]. Additionally, some have shown that increased triglyceride content impacts ventricular mass and cardiac work [47]. We believe that calorie restriction blocks metabolic reprogramming following cardiac hypertrophy induction by improved lipid synthesis/degradation or prevention of lipid oxidative damage. These result from decreased mitochondrial ROS generation and improved antioxidant enzyme levels (seen in Figure 1). Additionally, calorie restriction induces an alteration of the type of substrate used from carbohydrate metabolism to fat metabolism in the liver [48]. Whether that is true for the cardiac tissue remains to be tested. Knowing that cardiac tissue preferentially uses fatty acids as substrates, one could speculate that calorie restriction would induce a type of metabolic flexibility that renders cardiac tissue less sensitive to the so-called metabolic switch [49]. This possibility needs to be further tested.

This study also provides relevant insights into the link between mitochondrial function, oxidative stress, and cardiac hypertrophy. Mitochondrial ROS potentially damage cardiac components, leading to impairment of the organ. Indeed, ROS can modify DNA, proteins, and lipids. In particular, we show that calorie restriction improved succinate-supported mitochondrial respiration, avoided high mitochondrial ROS production, and antioxidant (MnSOD, GPX) repression during isoproterenol-induced cardiac hypertrophy. We believe mitochondrial protection is a central part of calorie restriction-induced anti-hypertrophic effects. Indeed, the literature has already demonstrated taht calorie restriction is a protective non-pharmacological intervention against cardiovascular disease [14]. Calorie restriction reduces blood pressure (a known risk factor for the development of cardiac hypertrophy in humans [18]) and preserves diastolic function [17]. Finally, we and others have demonstrated that calorie restriction is a powerful intervention able to block cardiac hypertrophy [15,16,19]. Mitochondria, together with peroxisomes, are the main sites for lipid degradation. The chemical energy stored in fatty acids and glucose is transformed into mechanical and electrical energy, favoring heart function. The majority of ATP in cardiac muscle is provided by mitochondria and used for cardiac muscle contraction and electrical activity [50]. Mitochondria are also the main intracellular source of ROS. Preservation of redox balance is important to control several heart functions such as excitability and contractile work [51]. Here, we show that hypertrophic cardiac mitochondria produce more ROS, accompanied by changes in cellular lipids. Additionally, we show that these changes are reversed by calorie restriction. Consistent with these observations, we have previously shown that calorie restriction blocks cardiac hypertrophy by decreasing cardiac ROS formation and by enhancing superoxide dismutase, catalase, and glutathione peroxidase [51]. Indeed, several lines of evidence associate mitochondrial oxidative stress with the development of cardiac hypertrophy/heart failure [4]. A possibility exists that many of the above-reported changes in lipids could be explained by the ability of ROS to promote cardiac lipid peroxidation, especially targeting the oxidation of polyunsaturated fatty acids. Furthermore, our results show that calorie restriction could lower hypertrophic-induced mitochondrial ROS, leading to anti-hypertrophic effects seen in markers of cardiac hypertrophy and in the lipidomic analysis. This study reveals calorie restriction as a resource to investigate biomarkers and potentially target cardiac hypertrophy-induced lipidomic, redox, and mitochondrial remodeling.

## CRediT authorship contribution statement

Study concept and design: Heberty T Facundo; Cicera Edna Barbosa David; Data acquisition: Cicera Edna Barbosa David, Pedro Lourenzo Oliveira Cunha, Yuana Ivia Ponte Viana, Anna Lídia Nunes Varela, Aline Maria Brito Lucas; Data analysis: Cicera Edna Barbosa David, Marcos Yukio Yoshinaga; Adriano B. Chaves Filho, Sayuri Miyamoto. Heberty T Facundo and Cicera Edna Barbosa David wrote the first draft of the paper, which was critically reviewed by Alicia Juliana Kowaltowski. All authors read and approved the final manuscript.

## Acknowledgments

The authors acknowledge the technical assistance of Camille C. da Silva and Antônio F. R. Santos.

---

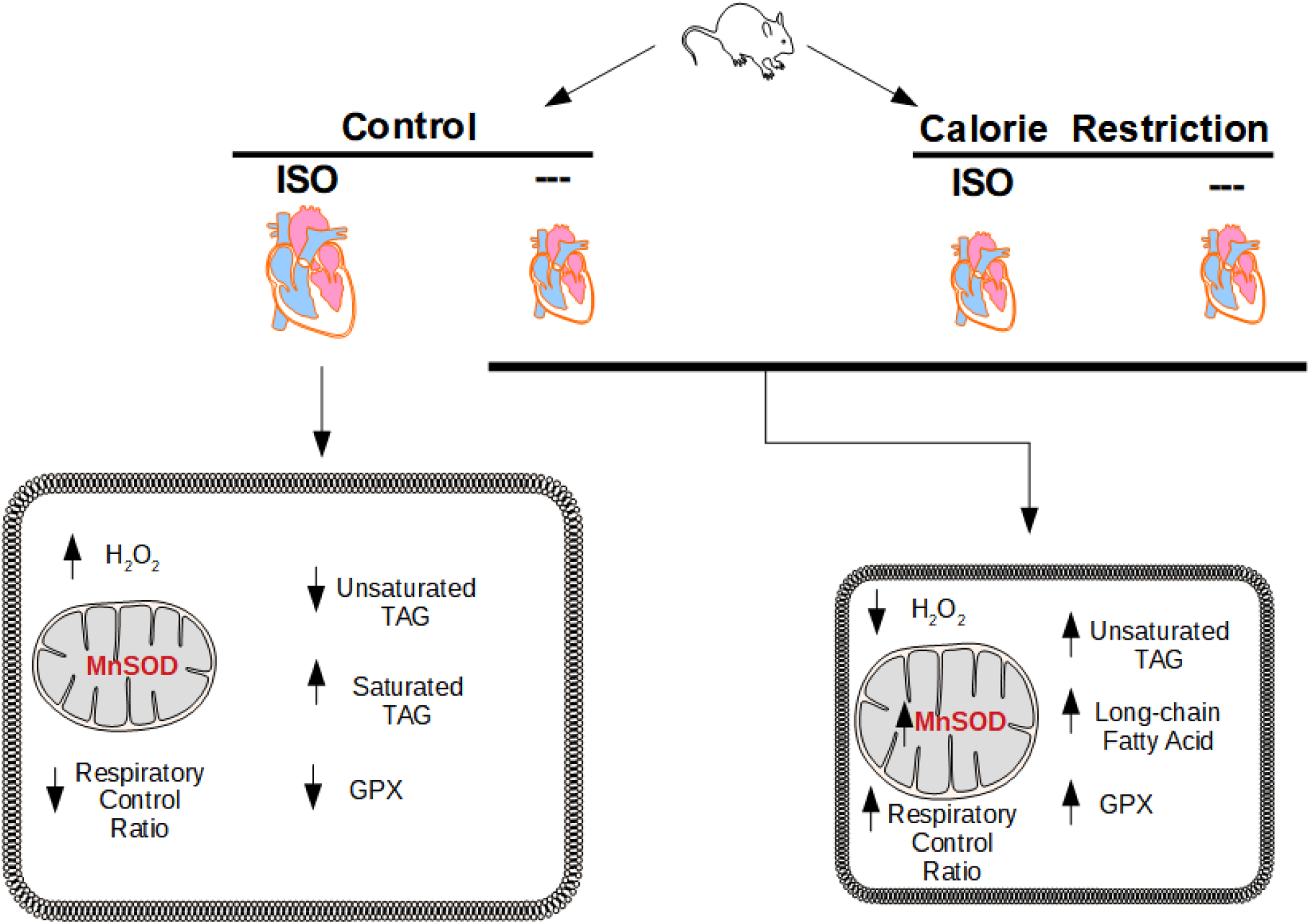

**Fig. S1.** Lipidome profile in hearts of isoproterenol-treated mice under calorie restriction. (A) Number of identified lipid molecular species per lipid subclass. (B) Concentration of the most abundant lipid subclasses. (C) Concentration of the less abundant lipid subclasses. Concentrations (ng/mg of tissue) are shown as mean ± standard error of mean (SEM). All groups were compared by one-way ANOVA followed by Tukey’s post-hoc (p<0.05). No significant alterations were observed.

**Fig. S2.** Volcano plot of other pairwise comparisons. (A) Volcano plot of the pairwise CRC versus ALC group comparison. (B) Volcano plot of the pairwise CRC_i versus AL_i group comparison. (C) Volcano plot of the pairwise CRC_i versus ALC group comparison. (D) Volcano plot of the pairwise AL_i versus CRC group comparison. Volcano plots are represented by log2 (fold change) plotted against-log_10_ (p value). Statistical significance was evaluated by t-tests (p < 0.05) and the fold change was set to >1.0.

**Fig. S3.** Veen diagram showing specific and common alterations in the lipidome of the experimental groups. (A) Veen diagram showing exclusive and common alterations (t-test, p < 0.05) between the groups: AL_i vs ALC, AL_i vs RCC_i and AL_i vs RCC. (B) Veen diagram showing exclusive and common alterations (t-test, p < 0.05) between the groups: RCC_i vs RCC, RCC_i vs ALC and ALC vs RCC.

