## Supplementary figures and images for "Calorie Restriction Maintains Mitochondrial Function and Redox Balance Avoiding Lipidomic Reprogramming during Isoproterenol-Induced Cardiac Hypertrophy"

### Figure S1

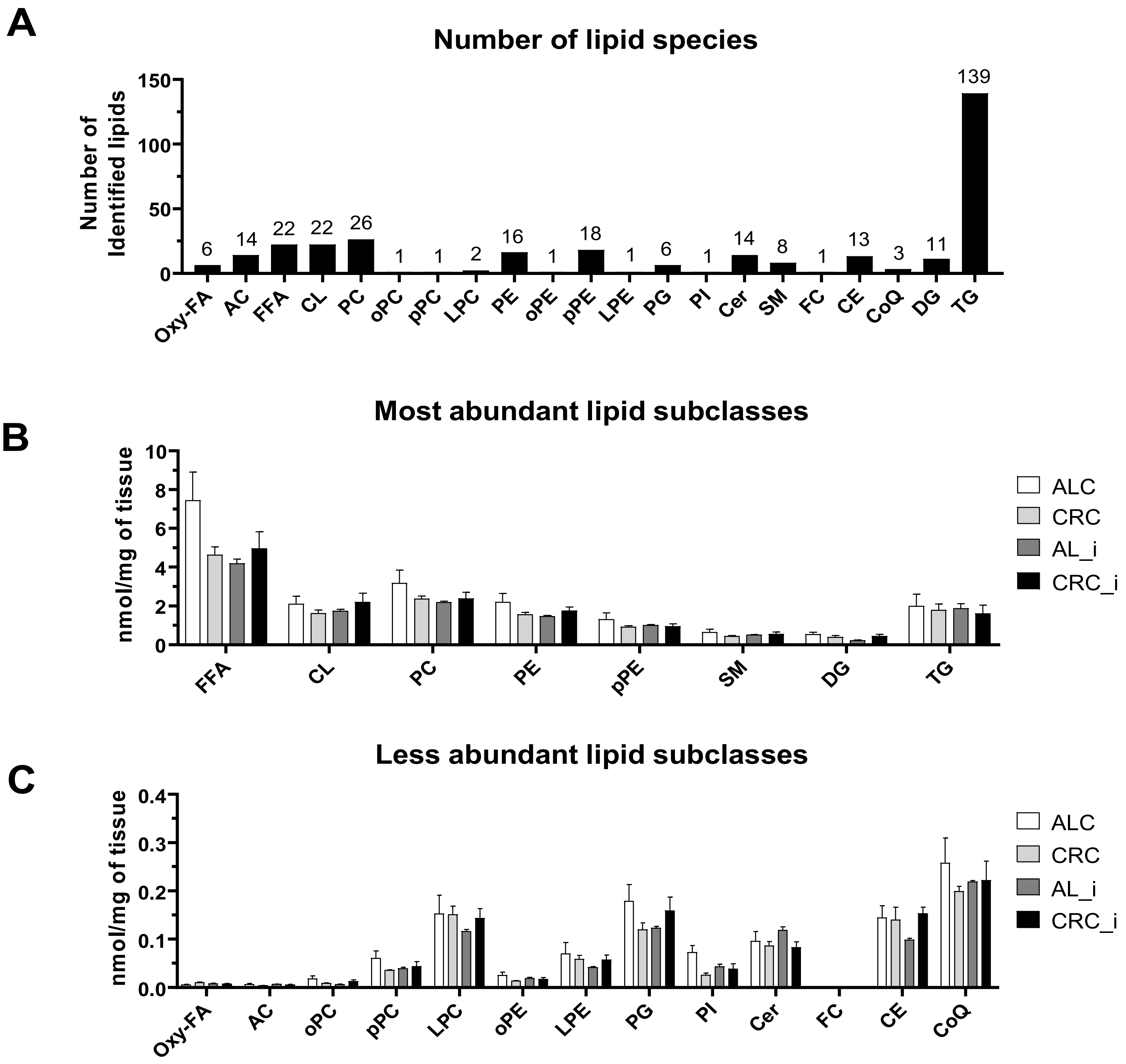

### Figure S2

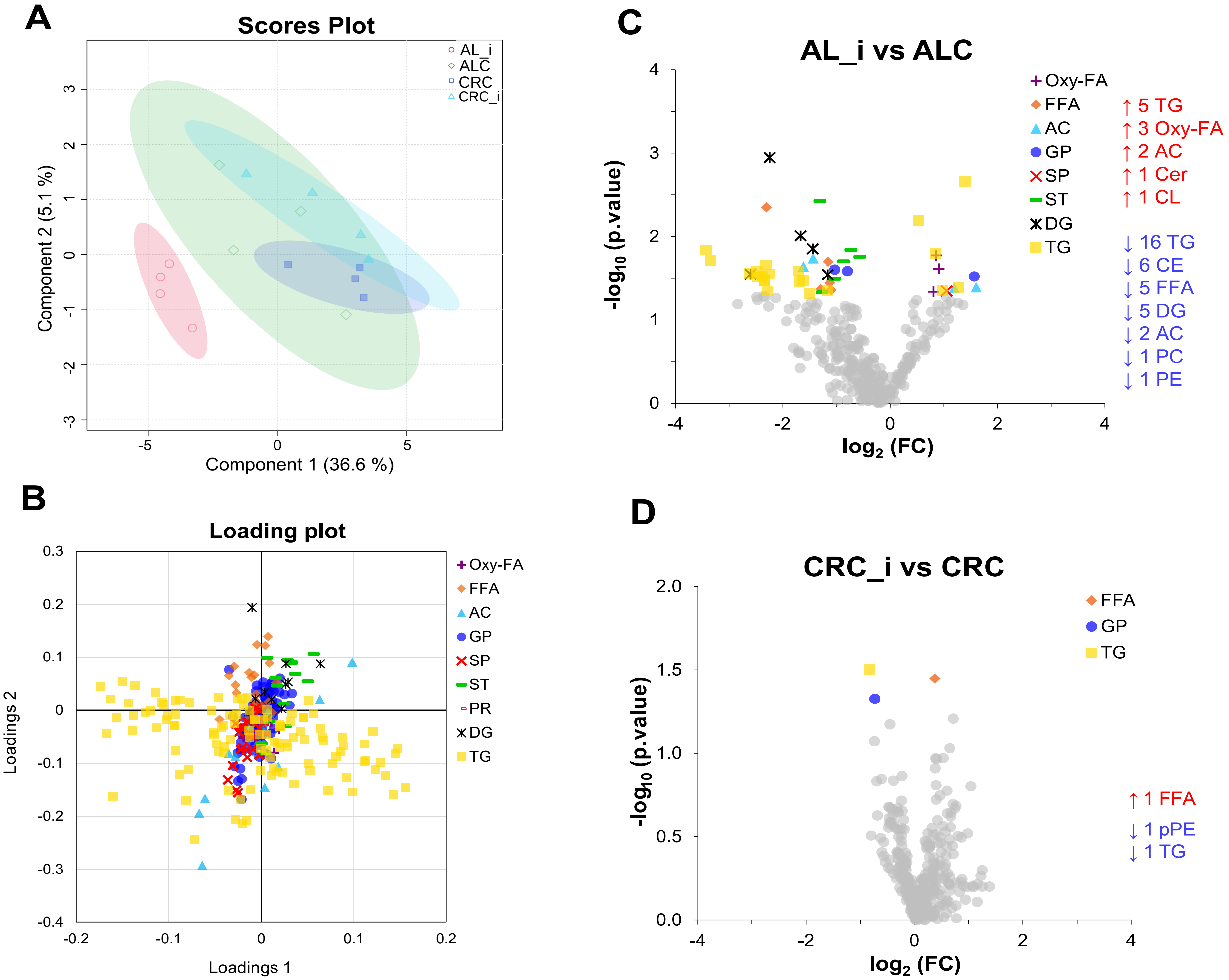

### Figure S3

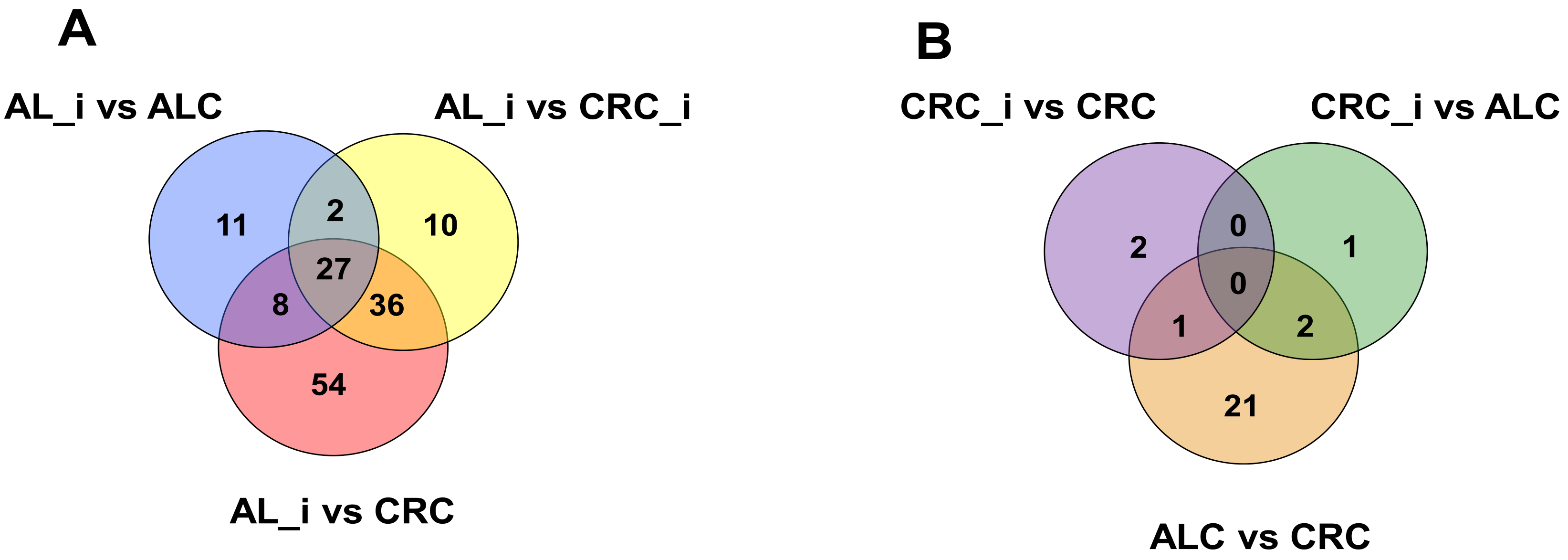
